# The Knock-In Atlas: A web resource for targeted protein trap by CRISPR/Cas9 in human and mouse cell lines

**DOI:** 10.1101/2025.02.19.638564

**Authors:** Yuma Hanai, Patrick Louis Lagman Hilario, Yuriko Shiraishi, Nobuyasu Yoshida, Suzuna Murakami, Yuji Shimizu, Norisuke Kano, Minami Kojima, Kokoro Murai, Taro Kawai, Katsutomo Okamura

## Abstract

Various cell engineering techniques have been developed by leveraging the CRISPR-Cas9 technology, but large-scale resources for targeted gene knock-in are still limited. Here we introduce the Knock-in Atlas, a web resource for gene tagging by fluorescent proteins by inserting artificial exons in target gene introns. To produce knock-in cells efficiently and reproducibly, we carefully chose and catalogued guide RNAs (gRNAs) for targeting genes in the human and mouse genomes by taking the gRNA efficacy scores and protein structures around the insertion sites into account. As of August 2025, we have characterized knock-in cell lines for 350 proteins, with a focus on RNA binding proteins, by flow cytometry and confocal microscopy. The transfection and flow cytometry protocols were optimized for several cell lines including HEK293T, eHAP1, HeLa, THP-1, Neuro2a, mouse embryonic fibroblast (MEF) and mouse embryonic stem cell (mESC). A website has been launched to organize the results of initial characterization including flow cytometry data after transfection, confocal microscopy, and western blot results for the genes for which knock-in HEK293T cell lines were already made. The site also provides a database to organize the information of pre-designed gRNAs for the human and mouse genomes. <https://rnabio.naist.jp/atlas/>.

## Introduction

Gene knock-in technologies have advanced to support a range of molecular biology applications, including gene sequence manipulation and tagging of target genes with epitopes, fluorescent proteins, or enzymes using various CRISPR/Cas9-based methods (1). For tagging proteins with fluorescent proteins or epitope tags, a large scale knock-in cell library was developed by using the homology-directed repair (HDR) after introducing double-strand breaks near the tag-insertion sites, producing knock-in HEK293T lines for >1, 000 protein targets (2). While this serves as an invaluable resource, not many laboratories would have the capability of targeting even hundreds of proteins in this way because this method requires construction of costly donor DNA unique to each target in addition to plasmids for gene specific guide RNA (gRNA) expression. The donor plasmids would have to be remade for all genes when using a different tag as well.

Methods utilizing non-homologous end-joining (NHEJ) hold the potential for cost-efficient large-scale cell line production because a single donor sequence could be used for insertions at various sites unlike those depending on HDR (3–5). The issues that arise from the frequent introduction of mutations at junction by NHEJ could be circumvented by inserting a synthetic donor, where an artificial exon encoding a tag protein sequence is flanked by artificial splice acceptor and donor sequences, in the target gene intron to produce in-frame fusion proteins (6–8). Because the mutations would not affect splicing patterns unless deletions reach the splice donor or acceptor sequences or insertions accidently create a new splice site. Indeed, this principle was used to produce a large scale knock-in cell library in a recent paper (7).

Despite the available techniques and resources, there remain challenges in creating large sets of knock-in cell lines such as: 1) Setting up knock-in protocols optimized for different cell lines, 2) Designing effective gRNAs, 3) Selecting tagging sites to avoid disrupting protein structures, 4) Selecting genes that are expressed at reasonable levels.

Here, we provide a tool kit for NHEJ knock-in wherein a large set of gRNAs have been designed against human and mouse introns on a genome-wide scale while avoiding introns flanked by exons encoding protein domains with predictable rigid structures. A database of pre-designed gRNAs that is searchable by target gene symbols has been developed and each entry is associated with the TPM values of the host gene in common cell lines and the

AlphaFold2 pLDDT scores of the 3 amino acids flanking the predicted tag-insertion site. As an initial set, we release 429 knock-in HEK293T lines in which localization of the tagged proteins were analyzed using the Venus knock-in donor, and the gRNA information and initial characterization data of knock-in cells have been made publicly available.

## Materials and Methods

### Plasmids

To construct the donor plasmid, the Venus sequence was amplified from a plasmid (Gift from the Hiroshi Itoh lab at Nara Institute of Science and Technology) using the Venus F and R primers (Supplementary Table 1). The sequence encoding the linker amino acid sequence of GSGGSGGGS, and the splice acceptor/donor sequence and the gRNA target cleavage sites was synthesized as oligo DNA (Fasmac) and they were combined together with the Venus amplicon and the pUC19 plasmid (Supplementary Table 1) using the In-Fusion HD Cloning Kit (Takara Bio). Three plasmids were produced for different intron phases by using pairs of oligos shown in Supplementary Table 1. In addition, three plasmids were modified to convert Venus to mVenus by using the mVenus mutation F and R primers shown in Supplementary Table 1. The mVenus version of donor DNA was used for producing all knock-in lines except HNRNPA1 knock-ins and those done in mESCs, MEFs and THP-1. The examination of Venus positivity rates was done using donor DNA with the original Venus, not the mVenus. To modify donor DNA cassette with different chemical modifications, the Venus cassette of the plasmid was amplified using Cassette F and R primers (Supplementary Table 1). To obtain the gRNA expression plasmid against target intron shown in Supplementary Table 2, the gRNA sequence predicted by CHOPCHOP (9) was synthesized with the sense strand added with 5’-CACC (or 5’-CACCG when the 5’ nucleotide was an H) and the antisense strand added with AAAC-3’ and inserted in pSpCas9(BB)-2A-miRFP670 (Addgene, #91854) or pSpCas9(BB)-2A-Puro (PX459) V2.0 (Addgene, #62988) digested with Bbs I (NEB) according to a previous study (10). The gRNA sequence against the donor plasmid referred as previous study (3–5).

### Oligonucleotide labeling

Primers with 2PS bond and C6-amine were synthesized as oligo DNA (Fasmac) shown in Supplementary Table 1. 5’-chemical modifications with PEG10 NHS ester (BROADPHARM) or Biotin NHS ester (Sigma-Aldrich) for C6 amine were performed following to a previous study (11). In brief, 1 mM NHS esters were incubated with 10 μM of primer with C6 amine group in 1×Borate Buffer (Thermo Fisher Scientific) overnight at room temperature. The primers were purified with Micro Bio-Spin P-30 Gel Columns, Tris Buffer (Bio-Rad). 10 pmol primers were resolved on a 15% SequaGel Sequencing System (National Diagnostics) and the gel was stained with SYBR Gold (Invitrogen). To obtain modified PCR product, the target region was amplified by PCR using KOD FX Neo (TOYOBO). The gel showed the efficient addition of each moiety (C6, C6-PEG10 and C6-Biotin) was verified by altered gel mobility (Figure S1A), and PCR products were readily amplified with modified primers (Figure S1B).

### Establishment of knock-in cell lines

HEK293T, eHAP1, HeLa, Neuro2a or mESC were grown on 6-well or 24-well plates coated with Cellmatrix Type I-C (Nitta Gelatin) or 0.01-0.1% Gelatin and transfected with plasmids or PCR fragments at the ratio of gRNA expression plasmid : Venus donor = 0.076 pmol : 0.064 pmol, using the FuGENE HD Transfection Reagent (Promega) or PEI MAX (PSI) according to the manufacturer’s instructions, and FBS (Final concentration 20%) was added to the medium 2 hours after the transfection. MEF or THP-1 cells were transfected with plasmids at the ratio of gRNA expression plasmid : Venus donor = 0.342 pmol : 0.429 pmol, using electroporation using the Neon Transfection System (Life Technologies) according to the instructions. Cells were resuspended in 1×PBS (-) in the case of detection Venus positivity rates, or Sorting Buffer (1×PBS (-), 1 mM EDTA, 25 mM HEPES-KOH, 1% heat inactivated FBS) in the case of establishment of knock-in cell lines and transferred to 5 mL Round-Bottom Polystyrene Test Tubes with Cell Strainer Cap (Falcon), and Venus positive cells were detected using CytoFLEX S. The cells were sorted onto plates that were coated with Cellmatrix Type I-C (Nitta Gelatin) or 0.01-0.1% Gelatin using Cell Sorter MA900 (SONY) or FACS Aria Special Order (BD). After the Venus positive populations were expanded, single Venus positive cells were sorted against onto 96-well plate coated with Cellmatrix Type I-C (Nitta Gelatin) to establish knock-in clones derived from single cells. Acquired FCS files were processed by using flowCore package (12) for visualization on the database.

### Organization of representative intron annotation

Matched Annotation from NCBI and EMBL-EBI (MANE v.1.0) (13) for human and RefSeq Select (14) from (GCF_000001635.27) for mouse were used to determine the representative transcript isoform set. Then, introns in CDSs that intersected with the start or stop codon were excluded from the analysis. The intron phase information was collected for 171, 806 introns in 16, 993 genes (human) 174, 285 introns in 17, 715 genes (mouse) in total (Supplementary Table 3). The percentages of intron phase usages were calculated by dividing the number of introns in each phase by the number of total introns in CDSs. The GFF files containing the representative transcript coordinates were generated and sorted by using GFF3sort (15), then, the TBI index files were generated by using Tabix (16) for visualization on UCSC genome browser.

### Design of gRNA sequences

CHOPCHOP was locally installed under the recommended environment (Python = 2.7.18, Numpy = 1.16.6, Scipy = 1.2.1, pandas = 0.24.02, Biopython = 1.74, argparse = 1.4.0, scikit-learn = 0.18.1)(9). gRNAs were designed by CHOPCHOP with the following options: -T 1 -f GN -M NGG -backbone AGGCTAGTCCGT -scoringMethod DOENCH_2016 -G hg38 or mm39 -Target[chr]:Intron[N]_start site_ - Intron[N]_end site_. The input genomic coordinates were prepared by modifying the annotated intron coordinates. The 75 nt from the exon/intron junction were excluded. In addition, introns longer than 40, 000 nt were split into <=40, 000 nt inputs for efficient CHOPCHOP runs. The cleavage efficiencies of gRNAs were predicted based on the machine-learning model developed by Doench et al., (17). gRNAs that have potential off-targets with no mismatch were omitted, and those with GC content within the 40-70% range and self-complementarity less than 1 nt were selected. Finally, the three top ranked gRNAs according to the Doench score are shown on the database. In total, this pipeline produced gRNA sequences against human 142, 862 introns in 16, 362 human genes and 149, 561 introns in 17, 398 mouse genes (Supplementary Table 4 and 5). The Bed files which contain the gRNA coordinates were generated and sorted by using BEDtools (18), then, bed files were converted into BigBed files by using bedToBigBed (19) for visualization on UCSC genome browser.

### Mapping of pLDDT scores on the human and mouse genomes

PDB files were downloaded from the AlphaFold Protein Structure Database (20), among which 17, 485 human proteins and 14, 868 mouse proteins matched the MANE v.1.0 or mouse RefSeq Select transcripts. In addition, 15 human proteins (ANXA7, DSP, GSPT1, HNRNPA2B1, HNRNPAB, HNRNPC, HNRNPK, PABPC4, PCBP2, PRRC2C, PTBP1, PTBP3, PUM1, PUM2, XRN1) that were from the MANE v.1.0 were individually predicted by AlphaFold2 (21). The predicted pLDDT scores were computationally mapped to the human and mouse genomes at the corresponding codons. In brief, pLDDT scores were extracted by using bio3d package (22) and corresponding amino acid coordinates were computationally calculated. Bed files which contain protein coordinate and corresponding pLDDT score were generated and sorted by using BEDtools (18), then, the bed files were converted into BigBed files by using bedToBigBed (19) for visualization on the UCSC genome browser. The average of pLDDT scores within the 3 amino acids surrounding the predicted tag insertion site was calculated for 139, 913 introns in 15, 071 human genes and 119, 380 introns in 13, 119 mouse genes (Supplementary Table 6).

### Website

The two links on the cell line table are to the experimental webpages for our data and the subcellular localization pages hosted by The Human Protein Atlas (23). UCSC genome browser (24) was embedded on the database to visualize the gRNA scores and pLDDT scores on representative transcript isoforms.

### Microscopy

Knock-in cells were grown on 96-well plates (ibidi or MATSUNAMI) coated with 0.1% gelatin in the case of mESCs or Cellmatrix Type I-C (Nitta Gelatin) for other cell lines. Cells were fixed with 4% formaldehyde for 15 minutes, and washed with 0.1% Triton-X for three times followed by a 10-minute permeablization with 0.2% Triton-X and washed with 0.1% Triton-X for three times. Cells were stained with Hoechst33342 for 5 minutes, and observed on ZEISS LSM980 (Carl Zeiss) with a 20× objective lens. The acquired CZI files were processed using ZEN blue edition (Carl Zeiss) and the images were exported to PNG files for visualization on the database.

### Splinkerette PCR and genotyping PCR

Splinkerette PCR was performed according to a protocol modified from a published method (25). eHAP1 knock-in cells were pelleted by a centrifugation at 1, 200 rpm at 4 degrees for 5 minutes and frozen at -80 degrees. 2 μg of genomic DNA extracted by NucleoSpin Tissue (Takara Bio) was digested by Mse I (NEB) at 37 degrees overnight and the enzymes were heat inactivated at 65 degrees for 20 minutes. The splinkerette adapters (Supplementary Table 7) were mixed and denatured at 95 degrees for 5 minutes, and annealed by cooling the samples at the rate of 0.1 degrees per second until the samples reaches 25 degrees. The annealed linkers were ligated to the digested genomic DNA was purified the FastGene Gel/PCR Extraction Kit (NIPPON Genetics). The target sequence was amplified by nested PCR using KOD FX Neo (TOYOBO) and the splinkerette and genotyping primers indicated in Supplementary Table 7. The PCR products were agarose-gel purified using the FastGene Gel/PCR Extraction Kit (NIPPON Genetics) and sequenced by the splinkerette sequencing primer (Supplementary Table 7).

### Western blotting

Cells were collected by a centrifugation at 1, 200 rpm at 4 degrees for 5 minutes, and lysed in 2×SDS Lysis Buffer (113.6 mM Tris-HCl pH 6.8, 2.3% SDS, 22.7% Glycerol, 90.9 mM DTT) or Lysis Buffer (50 mM Tris-HCl pH 7.4, 100 mM NaCl, 1% Igepal, 0.1% SDS, 0.5% Sodium deoxycholate). In the case of cell lysis by Lysis Buffer, total protein was quantified with the TaKaRa BCA Protein Assay Kit (Takara) and an appropriate amount of 2×SDS Lysis Buffer was added. After resolving the protein sample by SDS-PAGE, proteins were transferred onto Immobilon-P PVDF Membrane (Merck Millipore) in CAPS Buffer (20% MeOH, 0.05% SDS, 10 mM CAPS pH 11.0), and the membranes were blocked using Blocking Buffer (5% skim milk, 20 mM Tris-HCl pH 7.5, 138 mM NaCl, 0.1% Tween-20) for 30 minutes, and followed by washed with 1×TBST (20 mM Tris-HCl pH 7.5, 138 mM NaCl, 0.1% Tween-20). The membranes were incubated with the primary antibodies listed in Supplementary Table 8 for 1 hour at RT or 1-3 days at 4 degrees and washed five times with 1×TBST followed by incubation with the secondary antibodies (Supplementary Table 8) for 1 hour at room temperature. The membranes were washed again with 1×TBST for five times, and then incubated with ECL Primer Western Blotting Detection Reagent (Cytiva) for 5 minutes. The signals were detected by FUSION-Chemiluminescence Imaging System (Vilber-Lourmat). Acquired gel images were processed for visualization on the database. The membranes were stripped with Harsh Stripping Buffer (62.5 mM Tris-HCl pH 6.8, 2% SDS, 0.8% 2-mercaptoethanol), then incubation was repeated with another antibody.

### RT-PCR

RNA was extracted by TRIzol-LS (Termo Fisher Scientific) and cDNA was synthesized by SuperScript III First-strand Synthesis System (Invitrogen) and RT primer. The target was amplified by PCR using KOD FX Neo (TOYOBO) and RT-PCR primers indicated in Supplementary Table 7.

### Immunoprecipitation

Cells were collected by a centrifugation at 1, 200 rpm at 4 degrees for 5 minutes, and lysed in Lysis Buffer containing a Proteinase inhibitor cocktail (Roche or Wako), and incubated the sample at 4 degrees for 5 minutes. The samples were sonicated at Low settings for 5 minutes (30 sec ON, 30 sec OFF, 5 cycles) using Biruptor, and the supernatant was collected by a centrifugation at 15, 000 g at 4 degrees for 10 minutes. Total protein was quantified using the TaKaRa BCA Protein Assay Kit (Takara), and 150-200 μg of total protein was used for immunoprecipitation. 500 μL of the sample diluted with Dilution Buffer (10 mM Tris-HCl pH 7.4, 150 mM NaCl, 0.5 mM EDTA) was incubated with 5 μL of GFP-Trap Magnetic Agarose (Chromo Tek) with continuous rotation at 4 degrees for 1 hour. The beads were subsequently washed two times with High-Salt Wash Buffer (50 mM Tris-HCl pH 7.4, 1M NaCl, 1 mM EDTA, 1% Igepal, 0.1% SDS, 0.5% Sodium deoxycholate), 1 time with High-Salt Wash Buffer + Wash Buffer (20 mM Tris-HCl pH 7.4, 10 mM MgCl_2_, 5 mM NaCl, 0.2% Tween-20), then 3 times with Wash Buffer. The washed beads were resuspended in an appropriate amount of 2×SDS Lysis Buffer, and the mixtures were incubated at 98 degrees for 5 minutes. The supernatant was separated from beads on a magnetic tube rack. After resolving the protein sample by SDS-PAGE, gels were fixed with Fix solution (40% EtOH, 10% Acetic Acid) for 2 hours, followed by stained with Flamingo Fluorescent Protein Gel Stain (Bio-Rad). Western blotting was done as mentioned above.

## Results and Discussion

### Designing targeting donors

In order to produce a knock-in library, we first considered several different forms of donor DNA. For simple detection of knock-in events, we used the Venus fluorescent tag that is flanked by splice acceptor and donor sites (Figure 1A left). Some prior studies used split-fluorescent proteins (e.g. OpenCell), with which the insertion sequences could be much shorter (26). However, in this study, we used the full-length Venus-tag because testing the efficiency using such a tag was more suitable for our aim, which is to develop a useful resource that many researchers can adopt for their experimental settings because prior establishment of stable cell lines to express the β-sheet 1-10 would be required to obtain split fluorescent-protein tag knock-in lines. Such a prerequisite could become a burden in certain situations. On the other hand, we note that our gRNA library could be used in conjunction with a variety of tags including split fluorescent tags in theory.

**Figure 1.**
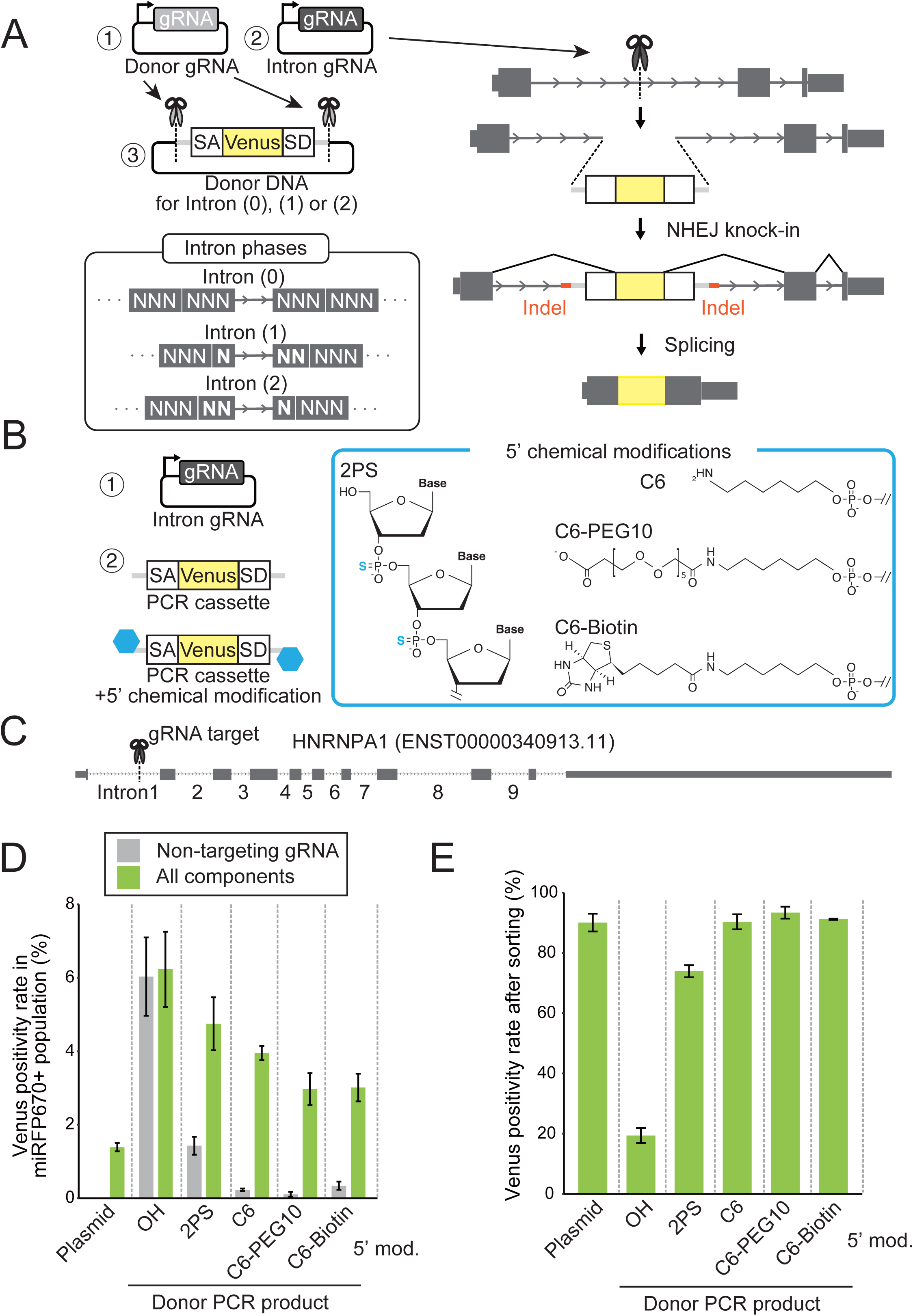
Evaluation of donor DNA in different forms. (A) Schematic of the strategy of the plasmid-based NHEJ knock-in. Three plasmids are simultaneously transfected: 1. gRNA expression plasmid to excise the Venus cassette from the donor plasmid, 2. gRNA expression plasmid to cleave the target intron, 3. Donor plasmid containing the Venus sequence flanked by the splice acceptor (SA) and splice donor (SD). Depending on the reading frame of the surrounding exons one of the three donor plasmids with the correct reading frame (Intron 0, 1 or 2) is used. (B) Utilization of modified PCR products in NHEJ knock-in. Instead of relying on the excision of the Venus cassette from the plasmid by CRISPR/Cas9 in cells, knock-in using donor DNA in the form of PCR products with different chemical modification at the 5’ end was examined. In addition to PCR products produced with regular 5’-hydroxyl DNA primers, those with 2PS (phosphorothioate linkages at the first two nucleotides), 5’-C6, 5’-C6-PEG10 or 5’-C6-Biotin modification were examined. (C) HNRNPA1 gene locus. A gRNA targeting HNRNPA1-Intron1 was used in experiments in panels (D) and (E). (D) Flow cytometry results of knock-in cells 2 days after transfection. The y-axis shows the percentage of Venus positive cells within the transfected cell population as judged by the presence of miRFP670 fluorescence derived from the gRNA expression plasmid backbone. The scatter plots used to calculate Venus positivity rates are shown in Figure S2A. Grey bars show the knock-in results with the non-targeting gRNA expression plasmid, gRNA expression plasmid against the donor sequence and the Venus donor (in the case of plasmids) or the PCR product. Green bars show the results with the complete set of gRNA expression plasmids and donor DNA. Data represent mean±SD (n = 3). (E) Flow cytometry results of knock-in cells at the second sorting. Cells after initial Venus positive sorting shown in green bars in (D) were grown, and the resulting populations were analyzed again by flow cytometry. The scatter plots used to calculate Venus positivity rates are shown in Figure S2B. Data represent mean±SD.

Introns belong to one of three phases, intron-(0), intron-(1) and intron-(2), where introns are located before the 1^st^ nucleotide, between the 1^st^ and the 2^nd^ and between the 2^nd^ and the 3^rd^ nucleotides in a codon, respectively. In the human and mouse genomes, the three intron phases, intron-(0), intron-(1) and intron-(2) are present at a ∼5:3:2 ratio in the CDS (Figure S2). To account for the reading frames in different intron phases, we prepared donor DNA with 0, 1 or 2 nt insertions (Figure 1A, Supplementary Table 1). Such constructs have been previously successfully utilized for NHEJ-mediated gene tagging by using donor plasmid DNA that was designed to be cleaved by a gRNA to release the exon cassette in cells (6, 8, 27). To test other forms of donor DNA, we used PCR products with various 5’ modifications to minimize immunostimulation and exonuclease-mediated degradation, and compared them with the conventional plasmid-based method (Figure 1B).

We tested the four forms of donor DNA to insert them in intron1 of the HNRNPA1 locus in HEK293T cells (Figure 1C). As a positive control, we transfected cells with the donor DNA plasmid along with a plasmid expressing a gRNA to release donor DNA, in addition to a plasmid for the gRNA against HNRNPA1-Intron1 similar to the previously established strategy (6). This method achieved a Venus positivity rate of ∼1.5%, indicating successful knock-in at a detectable frequency (Figure 1D and S3A). When cells were transfected with the donor PCR product without 5’ modifications, we observed about 5% of “Venus positive” cells even when the Cas9-gRNA plasmid was omitted, but we assumed these cells to be damaged cells which often produce autofluorescence. Chemical modifications either in the 5’ region at the phosphodiester bridges (phosphorothioate bridges at first 2 nucleotides: 2PS) or the 5’ modifications linked with a C6 chain resulted in partial (2PS) or almost complete (5’ modifications) suppression of the appearance of presumable autofluorescent cells (Figure 1D, Figure S3A), similar to the observations seen with HDR-mediated knock-in experiments 1-2 days after transfection with PCR products (11). Various Venus positivity rates (3-4%) were observed with the PCR products with the chemically modified primers at the first sorting 1 weeks after transfection, and became > 86% Venus positive populations after the amplification of sorted cells, except the sample transfected with the unmodified PCR product (Figure 1E, Figure S3B). Knock-in using the unmodified PCR products failed to produce a population with a high percentage of Venus positive cells after expanding sorted cells. This might be attributable to the high percentage of presumable autofluorescent cells that interfered with the enrichment of Venus positive cells at the initial cell sorting. While the mechanism behind this phenomenon remains unclear, our results indicated that 5’ modifications, especially those with C6 or other additional modifications could be used as donor molecules, which may be especially beneficial when working with cell lines whose transfection efficiency is highly sensitive to the amount of DNA. The use of PCR products allows us not only to eliminate the unnecessary plasmid backbone of the donor, but also to omit gRNA against the donor plasmid.

We tested if this approach produces similar results in other cell lines. For this purpose, we used two human (eHAP1 and HeLa cells) and two mouse (Neuro2a and mESC) to perform similar experiments (Figure S4A). Interestingly, the relative Venus positivity rates with respect to the plasmid transfection were variable depending on the cell line, ranging from ∼1.74-fold enhancement in HeLa cells to very low in mESCs. Nevertheless, Venus positive cells could be collected by sorting (Figure S5). Therefore, the effectiveness of the method with modified PCR products appeared to be variable depending on the cell line or perhaps transfection conditions. Although we do not know the factors determining the effectiveness, we recommend the plasmid donor as the first choice as the knock-in worked relatively consistently with various conditions. C6-modified PCR products may be tested when plasmid donors produced poor results. Using the HNRNPA1-Intron1 gRNA, we tried our plasmid-based method on a human monocyte cell line THP-1 and mouse embryonic fibroblast (MEF) cells, which resulted in the successful production of Venus positive cells at the rate of ∼1% and ∼3%, respectively, further validating the application of the method in a broad range of cell types (Figure S4B).

Given the nature of the PCR donor as linear DNA, it was conceivable that such donors were prone to randomly integrate into the host genome, form concatemers, or become more susceptible to trimming by exonucleases. We first tested whether this strategy could produce frequent random insertions occurring independently of the gRNA-mediated genomic DNA cleavage events. We employed splinkerette PCR to amplify the sequences surrounding the inserted Venus sequence (25). We chose 5 independent HNRNPA1-Intron1 knock-in eHAP1 lines obtained with the C6 modified PCR-based method (Figure S6). The splinkerette PCR products from 5 out of 5 clones corresponded to the HNRNPA1-Intron1 sequence, however, an additional band was produced in one of the clones (Figure S6C), whose sequence indicated the insertion of Venus in the opposite direction, a potential drawback from our current NHEJ-based method. The actual random integration frequency remained unclear as only a small number of clones were tested. A previous study reported random integration of plasmid DNA and dsDNA that occurred 0.5-2.0% 21 days after transfection (28), and the absence of randomly inserted copies needs to be verified when the applications require knock-in lines free of extra insertions of the tag or the Cas9/gRNA expression cassette. We also note that these results do not exclude the possibility of off-target insertions when Cas9-mediated cleavage is not specific.

To test the integrity of the inserted tags, we also amplified the regions around the tag-insertion sites using primers against flanking sequences in cells with Venus insertions at intron 9 of the HNRNPA1 locus derived from the plasmid donor or the PCR donors with 5’ modifications (Figure S7A). Potential concatemer formation events were detected with both plasmid and PCR donors (Figure S7A and S7B, asterisks), and this phenomenon appears to occasionally occur broadly regardless of the target or the form of the tag (6). Although our results with a single locus and small number of clones did not allow us to estimate the frequency of concatemer insertion, users of NHEJ-mediated knock-in methods should be aware of this potential issue. When the sequences around the tag-insertions were analyzed, we observed indels in virtually all clones but these were akin to those seen when targeted-mutagenesis through NHEJ was induced by CRISPR/Cas9 (Figure S7C). Therefore, these results suggested that modified PCR fragments did not cause inaccurate knock-in more frequently than the plasmid-donor, but care should be taken considering these potential issues that may occur in any of the donor types discussed above.

Whether the tag is inserted at single or multiple alleles is an important factor for users, as some applications require all alleles to be tagged (for example when the protein is depleted by RNAi or the degron against the tag) and the presence of untagged alleles is preferred by some other applications. HEK293T is believed to be pseudotriploid, in which the majority of genes have three alleles although the copy number varies across the genome (29). At the HNRNPA1 locus in our culture, we consistently observed two alleles when the genomes of knock-in lines were analyzed and we assumed the locus has two copies. The PCR-based genotyping also allowed us to estimate the frequencies of single- and dual-allele tagging, and at least under this experimental condition, single-allele tagging was predominant (Figure S7). However, we note that the frequency can vary significantly depending on the target genes and experimental conditions.

Highly efficient splicing needs to occur to connect the inserted artificial exon and flanking exons to tag the target protein. To evaluate the splicing efficiency, homozygous knock-in clones for HNRNPA1 gene at introns 1-9 were prepared (Figure S8A). Despite the fact that these clones were verified as homozygous knock-in lines, extra bands were detected by Western blotting using an anti-HNRNPA1 antibody in clones with insertions at intron 3, 5 and 9 (Figure S8B). The shorter bands were not recognized by the GFP antibody, suggesting that the tag exon was skipped. Indeed, shorter bands near the wild-type HNRNPA1 cDNA size were seen in the clones with insertions at intron 3, 5 and 9, supporting the idea that the isoform skipping the tag-exon (Figure S8B). Closer inspection of the gRNA target locations in these clones revealed that the tags were inserted near the exon-intron junctions (51, 25, 80 nts away from the closest exon), suggesting that insertions near the splice sites should be avoided. Therefore, in our gRNA design pipeline, we excluded 75 nts from exons to avoid such splicing errors.

Insertion of fluorescent-tag potentially impacts on target protein functions. To test the effects of Venus-tag insertion on the protein functions, we used the homozygous Venus knock-in lines at all 9 introns (Figure S8A). A previous study showed that depletion of HNRNPA1 changes ∼2, 500 alternative splicing events, some of which were verified by RT-PCR assays (30). Using one of such well-characterized HNRNPA1 target genes NF1, we checked its splicing patterns in the 9 knock-in lines as a readout of HNRNPA1 activity because exon-skipping was more frequently seen with this gene when HNRNPA1 gene was knocked down. We found that the function of tagged proteins in regulating alternative splicing was impaired when the Venus tag was inserted near the RRM or RGG domains (introns 2-6) in contrast to the insertions in regions with lower pLDDT scores (Introns 1 and 7-9), which were relatively benign (Figure S8C). Therefore, this result validated our approach to avoid regions with high pLDDT scores.

### Designing gRNAs

Various factors affect the knock-in rates, among which gRNA design is presumably most important. Good gRNAs to efficiently produce high-quality knock-in cells would satisfy the following criteria: 1) The gRNA mediates robust and specific cleavage of genomic DNA at the target site, 2) The target intron is efficiently spliced, 3) The insertion of the tag does not strongly interfere with the target protein function. In addition, 4) Lowly expressed genes would typically be excluded because the fusion protein needs to be endogenously expressed at detectable levels for subsequent experiments. The following subsections explain how our pipeline designs gRNAs satisfying each of the criteria.

#### 1) Efficiency and specificity of gRNA cleavage

The cleavage efficiencies of gRNAs for all human and mouse introns of representative transcripts were predicted based on the machine-learning model developed by Doench et al., and deployed in the CHOPCHOP platform (9, 17) (Figure 2A). gRNAs that have potential off-targets with no mismatch were omitted, and those with GC contents within 40-70% were selected. Up to three gRNAs for each intron with the highest cleavage scores were listed.

**Figure 2.**
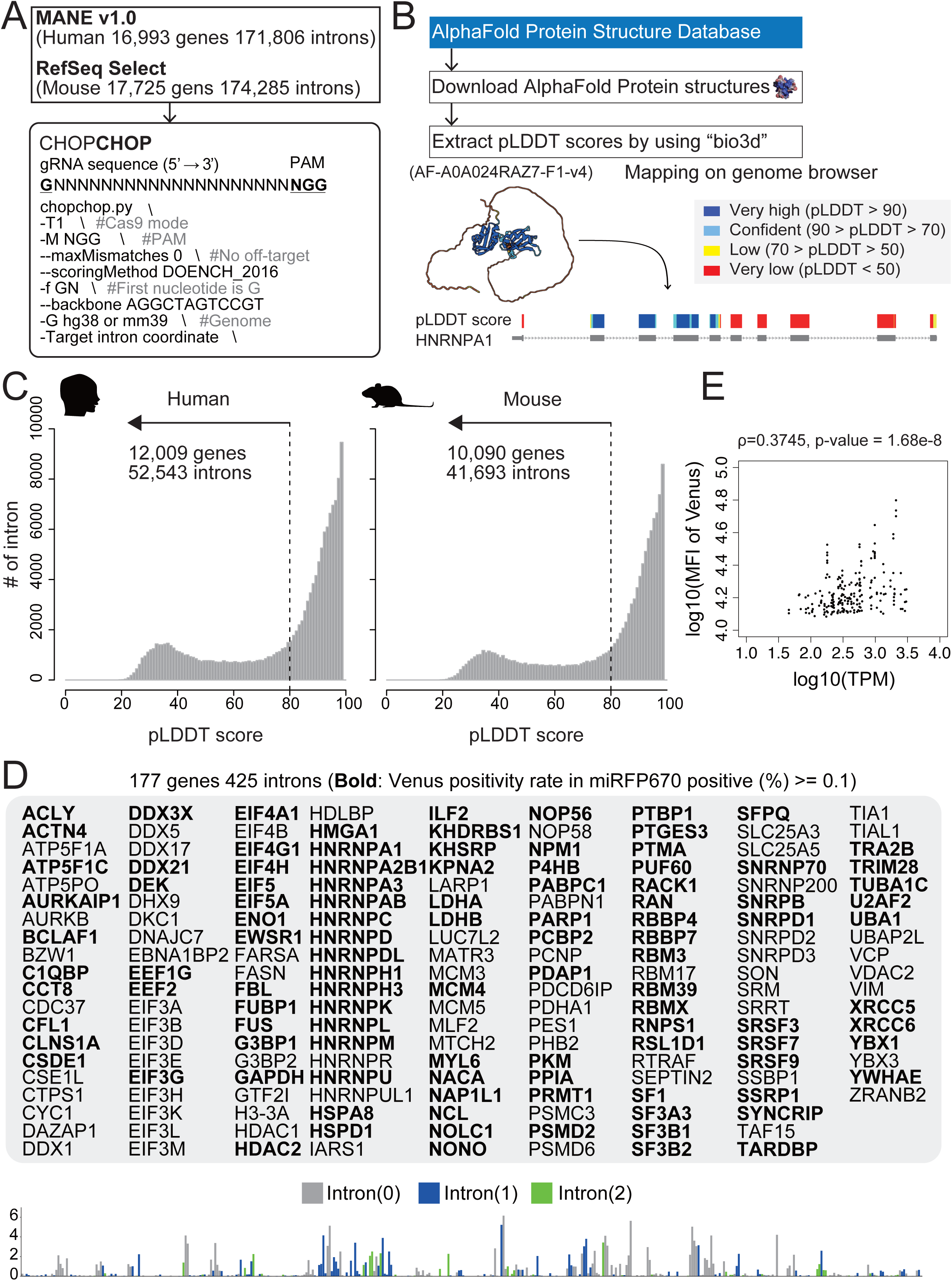
gRNA design pipeline. (A) Schematic of the gRNA prediction strategy. The representative transcripts were taken from the MANE v.1.0 and RefSeq Select references, and used the intron sequences for the CHOPCHOP prediction to obtain gRNA sequences. The parameters used are shown. (B) Visualization of pLDDT and gRNA scores on UCSC genome browser. The pLDDT scores were extracted from the AlphaFold Structure database by the bio3d package, and the scores were mapped to the genome at the corresponding codons. The pLDDT scores are color coded on the pLDDT score track, and the mapped scores at the human HNRNPA1 locus is shown as an example. (C) Distribution of mean of pLDDT scores within 3 amino acids surrounding the predicted tag insertion site. The averages of the pLDDT scores for the surrounding 3 amino acids were calculated for all human and mouse introns in the representative transcript set, and the histograms for human and mouse were drawn. The x and y-axes show pLDDT score and the number of introns in each bin, respectively. (D) Venus positivity rates of knock-in cells for 177 gens 425 introns. Flow cytometry of knock-in cells 2 days after transfection against 425 introns (intron-0: n = 183, intron-1: n = 148, intron-2: n = 94). The y-axis shows the percentage of Venus positive cells within the transfected cell population as judged by the presence of miRFP670 fluorescence derived from the gRNA expression plasmid backbone. 213 out of 425 shows more than 0.1% of Venus positivity rates. Data represent mean (n = 3) except for EIF3E-Intron8 (n = 2). (E) Correlation analysis between RNA expression levels in HEK293 cells and Venus positivity rates against each target gene. The observed Venus median fluorescent intensity (MFI) after transfection were plotted against the mRNA expression level of the corresponding gene in HEK293 cells in a published RNA-seq data (35). The correlation coefficient and p-values were calculated by spearman’s correlation analysis (n = 213).

#### 2) Selecting efficiently spliced introns

As mammalian genes produce multiple isoforms by alternative transcription start/termination and splicing, it is vitally important to select introns that are efficiently spliced for inserting our knock-in donor. A reasonable solution was to select a representative transcript isoform for each gene, which was catalogued as canonical isoforms in The Matched Annotation from NCBI and EMBL-EBI (MANE v.1.0) (13) and in RefSeq Select for human and mouse, respectively (14). In addition, we excluded the proximal and distal 75 nt of each intron from gRNA prediction, as they are within exon/intron junctions that often contain sequences important for splicing (31).

#### 3) Avoiding rigid protein structures

To avoid interfering with the functions of the target protein, a plausible way is to avoid ordered regions. The pLDDT scores produced by AlphaFold2 (21) are a promising genome-wide resource to predict such regions as tightly folded structures tend to be predicted more confidently (32). The predicted pLDDT scores were mapped to the human and mouse genomes at the codons encoding the corresponding protein sequence, and the average of pLDDT scores within the 3 amino acids surrounding the predicted tag insertion site was calculated (Figure 2B). The default pLDDT score threshold was set at 80 according to a previous study (32) but the users can modify the threshold using our interface (see below). With the cutoff of the pLDDT score at 80, there were 52, 534 and 41, 693 introns of 12, 009 and 10, 090 genes, in the human and mouse genomes, respectively (Figure 2C). In addition to these resources, average pLDDT scores within the 3 amino acids at the N- or C-terminus were calculated and reported in Supplementary Table 6 for those who are interested in inserting tags by other means such as HDR-mediated knock-in or plasmid-based overexpression.

#### 4) Expression level of the target gene

As the expression level of the tagged protein in knock-in cells is determined by the endogenous mechanisms via transcriptional and post-transcriptional regulatory elements, consideration of endogenous expression levels of its native counterpart protein is convenient for detecting fluorescently-tagged proteins after knock-in. For the practical application of knock-in cells, the tagged proteins need to be detectable by fluorescence from the tag (in the case of flow cytometry and cell imaging) or antibody-based detection (in the case of western blotting and immunostaining). As a proxy of the expression level, we used RNAseq TPM values. To understand the correlation between the TPM values and knock-in success rates, we measured Venus positivity rates of 425 knock-in targets, in which 213 gRNAs produced Venus positive cells at > 0.1% (Figure 2D). The Venus positivity rate showed a positive correlation (Spearman Rho= 0.3745, p-value = 1.68e-8) (Figure 2E). On our gRNA database, we set the threshold of normalized TPM at 45 as a default value, with which gRNAs would produce reporter lines with readily detectable fluorescence by flow cytometry (Figure 2D). We also note that some of the knock-in lines we successfully made had lower TPMs, as low as 20 (e.g. TIA1 gene). Therefore, users can adjust the threshold values using the slider depending on their needs (See below in the Web resource section). Currently, our interface includes RNAseq data from 4 human cell lines (HEK293, HeLa, eHAP1, and THP-1) and 2 mouse cell lines (MEF and mESC) (33–35).

### Web resource

A user-friendly interface has been launched to help individual researchers design good gRNAs for knock-in experiments. This website is mainly consisted of three pages, “Knock-in cell lines”, “Human gRNA database” and “Mouse gRNA database”. In the “Knock-in cell lines” page, the main table shows a list of knock-in lines, and contains an experimental summary to indicate whether flow cytometry data after knock-in transfection, fluorescent microscopy and Western blotting results are available (Figure 3A). The table contains two URL links, one of which prompts a new page to show our experimental results and the other to the webpage of The Human Protein Atlas of the corresponding gene (23) so the users can easily compare the results for the same gene in the two resources. Our experimental summary page shows the results with HEK293T and mESC including flow cytometry data of Venus intensity 2 or 4 days after transfection and after cell sorting, confocal microscopic images of bulk knock-in cells with Hoechst staining and Western blot gel pictures of knock-in lines derived from single cells using an anti-GFP antibody against the Venus tag (Figure 3B).

**Figure 3.**
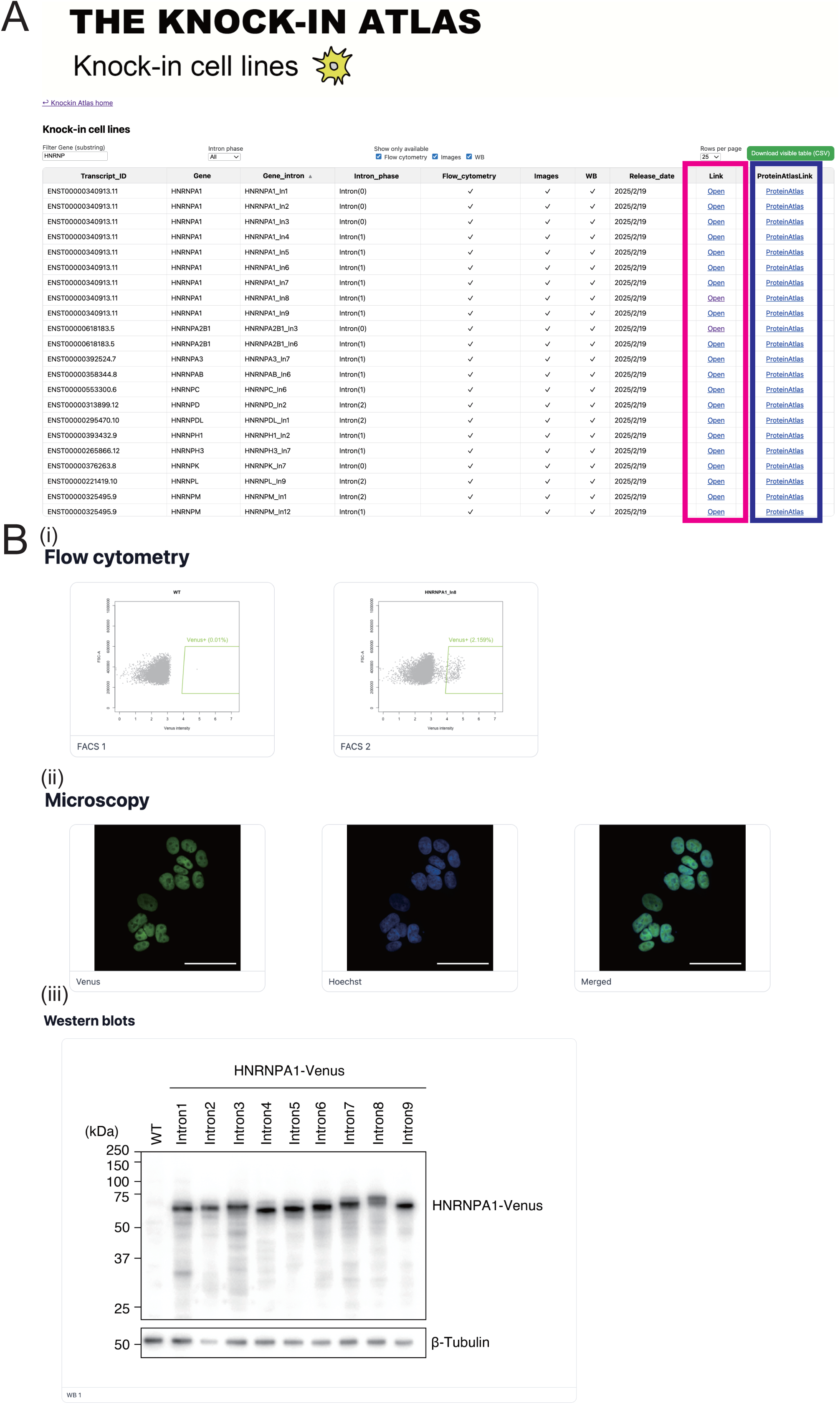
Experimental summary of knock-in cell lines. (A) Screenshot of the experimental summary page in the knock-in cell line database. The “Knock-in cell lines” page provides a table with last two columns showing URLs, one of which prompts the webpage summarizing our knock-in results (magenta rectangle) and the other the webpage of The Human Protein Atlas (dark blue rectangle). (B) The webpage of experimental summary for each target intron. The webpage contains 3 panes (i) Flow cytometry data of Venus positivity rates 2 days after transfection (ii) Fluorescent images of each bulk knock-in line (knock-in lines derived from single cells in the case of HNRNPA1) (iii) Western blots of each knock-in line derived from single cells.

The second and third pages show the pre-designed gRNA information for the human and mouse genomes, respectively. There are three parts with two sortable tables and a UCSC genome browser window (Figure 4). The first table is searchable by gene symbol or for representative gene functions (Figure 4A). When a row of the first table is selected, the user will see a list of introns within the CDS of the selected gene in the second table (Figure 4B) and the browser will show the selected gene locus (Figure 4C). The browser has a gene annotation track with a representative isoform for each gene, an AlphaFold2 pLDDT score track and a Pfam domain track (36). The second table shows a list of introns that satisfy the pLDDT score threshold (Set at < 80 by default, but can be adjusted using the slide bar) along with its intron phase and the pLDDT score in the adjacent 3 amino acids encoded by the surrounding exons. When a row of the second table is selected, up to 3 gRNAs with best gRNA cleavage scores for each intron will be shown in the third table (Figure 4D) and the browser will zoom into the selected intron. The third table shows the intron phase, gRNA strand, gRNA sequences and gRNA scores within the selected intron. For convenience, a download link for the csv file containing the information of the selected sRNAs is provided at the bottom of the page (Figure 4D).

**Figure 4.**
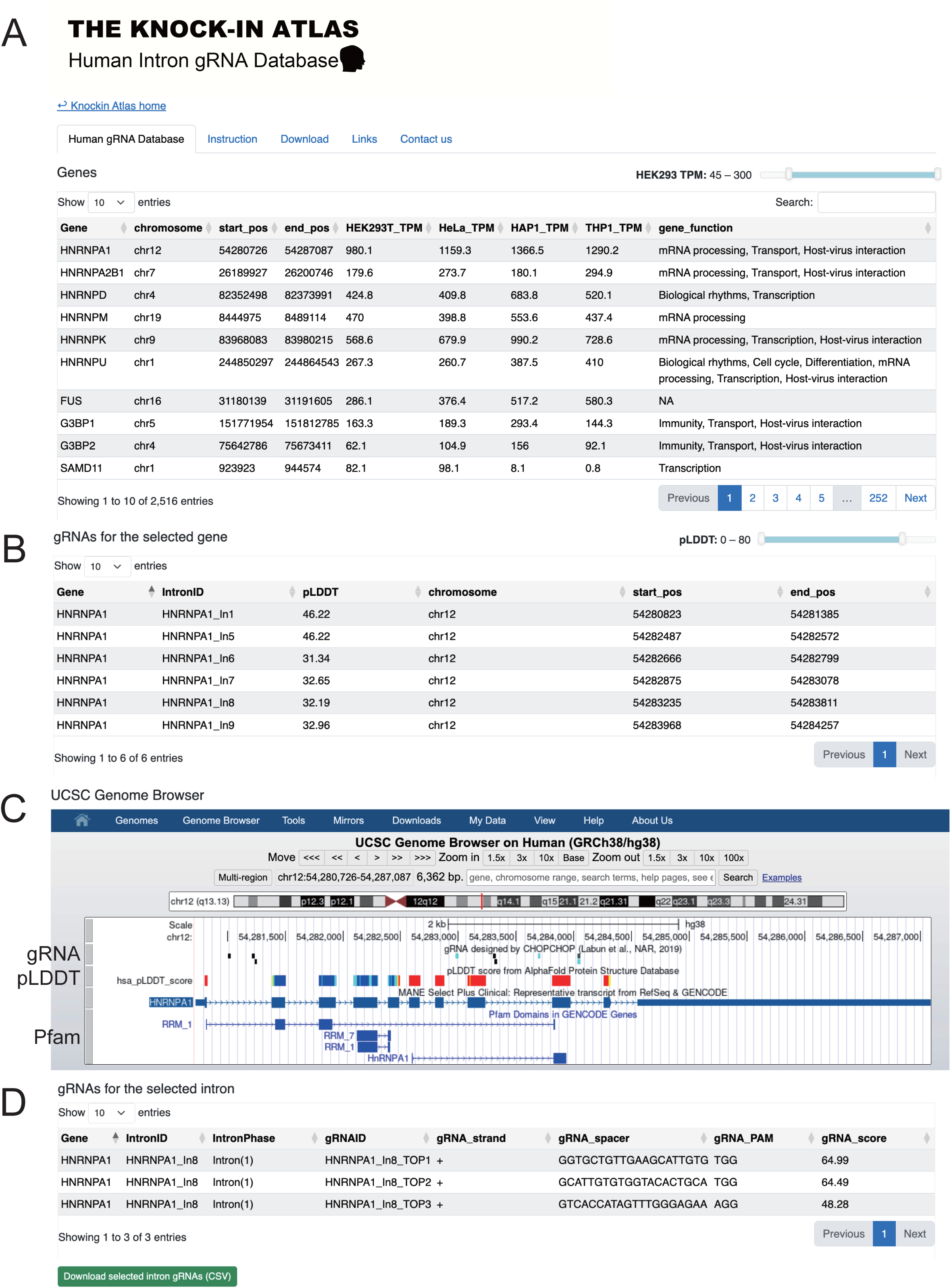
gRNA database for human and mouse. Screenshots of the gRNA database. Users can use the database to look for gRNAs for a target of interest using this database. There “Human gRNA database” and “Mouse gRNA database” pages and the human database is shown as an example. (A) The first table shows a list of genes, genomic coordinates and representative gene functions. (B) The second table shows a list of introns corresponding to the selected gene to show the intron phase, intron coordinates and pLDDT score. (C) The UCSC genome browser shows the selected gene locus or intron with three tracks, which display the representative transcript isoform, gRNA locations, pLDDT score and Pfam domains. (D) The third table shows a list of gRNA sequences corresponding to the selected intron along with the information of the intron phase, gRNA spacer sequence, gRNA PAM sequence and gRNA score.

### Estimated costs

To help users estimate the costs, below is the estimated cost breakdown following our current protocol. Plasmid construction and transfection/cell line production costed ∼ 6 USD (gRNA oligos, miniprep columns and DNA sequencing reagents) and ∼ 4 USD (cell culture medium, plates, transfection reagents) per target, respectively, excluding the costs for the usage of cell sorters and DNA sequencers.

Typically for 48 samples, the estimated hands on times are miniprep 2 hrs (2.5 min/sample), Sequencing 1 hr (0.8 min/sample), transfection 1 hr (0.8 min/sample), cell sorting 2-3 hrs (2.5-3.75 min/sample) and cell culture handling 1-2 hr (0.8-2.5 min/sample) and less trained personnel (undergrad interns even with non-biology major) has been able to handle most steps of the entire process in our current environment.

### Verification of results using public information

Among the 350 proteins analyzed in our project, protein localization information for 335 proteins was available at The Human Protein Atlas project (23) which allowed us to evaluate our knock-in lines. We counterchecked the localization of our target proteins against the Human Protein Atlas data, and 308 out of 335 knock-in lines showed roughly consistent localization results (Figure 5A). For example, our PTBP1 knock-in lines showed nuclear foci, which are known to locate at the perinucleolar compartment (PNC) that are often formed in malignant and transformed cells (37). Among the 27 proteins that showed localization patterns clearly different from The Human Protein Atlas information, the localization information of 21 of them (APP, ARPC2, CCT8, CDC123, CDKN2A, DEK, EFTUD2, EIF3L, HMGA1, HTATSF1, NPM1, PARP1, PSMC2, PSMF1, RAC1, RACK1, RBM3, SSRP1, STOML2, UBAP2L and XRCC6) was available at OpenCell and/or vpCells and showed similar patterns to ours (2, 7). We found additional independent evidence in the literature for 4 out of the 21 proteins by immunostaining or fusion protein expression (HMGA1, RACK1, SSRP1 and NPM1) (38–41). The remaining 6 proteins (BZW1, CFL1, COPS6, EIF5A, HEY1 and ILF2) were not present in either database, but we found literature information that supported the subcellular localization patterns determined in our results (42–47).

**Figure 5.**
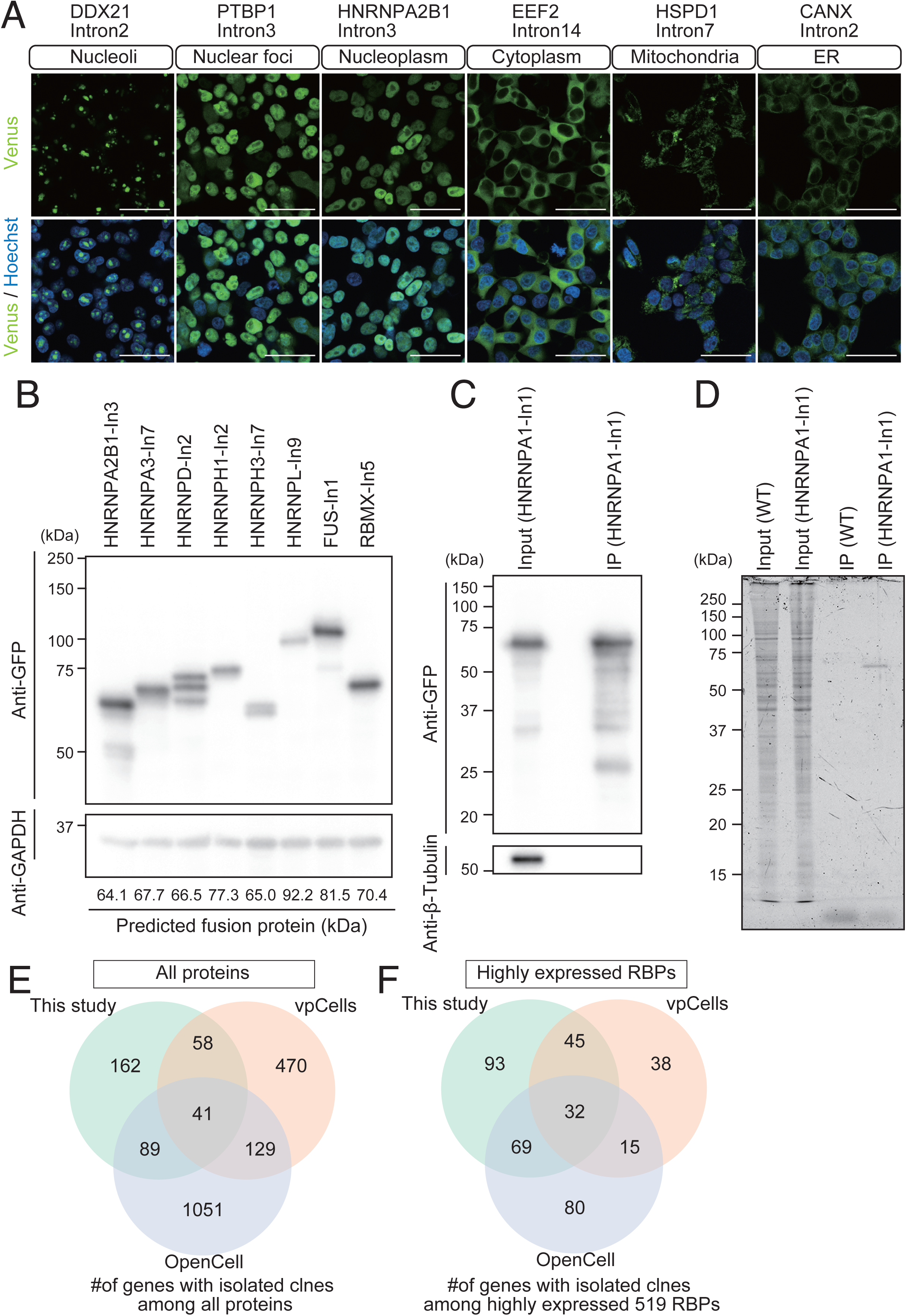
Examples of experiments using HEK293T knock-in lines. (A) Representative confocal microscope images of tagged proteins. Venus positive cells were enriched by a cell sorter and stained with Hoechst. The pictures were taken with a 20× objective lens on a confocal microscope, and scale bars are 50 μm. (B) Representative western blots of knock-in lines derived from single cells. Lysates from the indicated knock-in lines were resolved by 8% SDS-PAGE and bands were detected by an anti-GFP and anti-GAPDH antibodies sequentially. (C) Representative western blot of Venus-tagged HNRNPA1 protein after immunoprecipitation using anti-GFP VHH antibody. The HNRNPA1-Intron1 line derived from single cells was used and the protein was immunoprecipitated with the anti-GFP VHH antibody. Samples were resolved 12% SDS-PAGE and bands were detected by an anti-GFP and anti-β-Tubulin antibodies sequentially. (D) Purification of the Venus-tagged HNRNPA1 protein. Parental HEK293T was used as a negative control. The 12% SDS-PAGE gel was stained with flamingo staining solution and bands were detected by a fluorescence gel imager. (E) Overlaps with other knock-in resources. The Ven diagram shows the overlaps of HEK293T knock-in cell lines between our project (350 proteins), OpenCell (1, 310 proteins) and vpCells (698 proteins). (F) Overlaps of knock-in lines for highly expressed RBPs in HEK293T. 519 RBPs defined by RBPWorld are expressed at >160 TPM in HEK293T. The Ven diagram shows the overlaps of RBP knock-in cell lines between our project (239 RBPs), OpenCell (196 RBPs) and vpCells (130 RBPs).

The 15 proteins with no Human Protein Atlas entries were counterchecked with OpenCell and vpCells, and found 8 proteins (MYL6, TUBG1, COPZ1, KRTCAP2, ARF5, ARF3, COX5A, UQCRB) to be present in either or both of the databases. MYL6 showed somewhat distinct localization patters (cytoplasmic in our set, cytoplasmic filamentous structures in OpenCell, cytoplasmic-nuclear localization in vpCells) and can be due to different microscope acquisition conditions or the effects of the tag insertion. In other cases, the localization pattens were largely similar. For the 7 proteins (PPIA, EIF1, SLC25A39, ATP5MC1, ATP5MC3, TMEM106C, ATP5F1C) we did not find entries in either of the databases.

The results of one of such proteins (RACK1), which were initially done with the clone with the Venus insertion at its intron 3, were further verified by examining additional independent Venus knock-in lines with insertions at its introns 6 and 7 (Figure S9). Again, both Venus-RACK1 fusion proteins showed similar cytoplasmic localization. These results justify the use of our tagged cell line collection, at least for screening purposes. On the other hand, for more focused experiments using a smaller number of proteins of interest, verifying results with independent clones with multiple different tag-insertion sites would be recommended as is always the case in such tagging experiments.

The molecular weights of the 79 target proteins on Western blotting are also a good indicator of successful knock-in. 73 out of 79 knock-in lines derived from single cells showed roughly expected molecular sizes (Figure 5B), however, EWSR1, FUS, KHDRBS1, HNRNPU, NCL and SFPQ showed unexpectedly higher sizes. These are consistent with the results shown on the Abcam websites (Anti-EWSR1#ab133288, Anti-FUS#ab154141, Anti-KHDRBS1#ab76471, Anti-HNRNPU#ab172608, Anti-NCL#ab216006, Anti-SFPQ#ab38148), suggesting that the native proteins also show unexpected migration patterns on SDS-PAGE. Indeed, a published paper indicated that the native EWSR1 protein migrated at ∼90 kDa unlike its predicted size of 68 kDa, supporting that our Venus-EWSR1 protein was at the correct size (48).

The affinity of the Venus tag with various GFP antibodies provides opportunities for biochemical applications. We performed immunoprecipitation by the anti-GFP VHH antibody using randomly selected 7 knock-in lines derived from single cells and the target proteins were efficiently purified (Figure 5C, Figure S10). To determine the purity, we stained the gel with the Flamingo dye, and found that the target proteins were purified to homogeneity (Figure 5D). All the 7 immunoprecipitated proteins were detectable by Western blotting (Figure 5C, Figure S10). Therefore, our knock-in resource provides an attractive alternative to immunoprecipitation by antibodies against the native protein, especially when good IP-grade antibodies are not available for the protein of interest, or the user needs to maximize the cost-performance to conduct multiplexed experiments against a large number of target proteins.

### Overlaps with the existing databases

As parallel efforts, two large-scale knock-in cell line libraries have been published using HDR (Cho et al., 2022) or intronic NHEJ (Reicher et al., 2024) based on HEK293T cells, with 1, 310 and 698 proteins currently targeted, respectively as of August 2025. Although our microscopy results are currently only for 350 target proteins, about a half of them (162 targets) were unique to our set (Figure 5E). Among 519 highly expressed (>160 TPM) RBPs defined by RBPworld (2, 49), 196 and 130 RBPs were present in OpenCell and vpCells, respectively, making their union to have 279 knock-in lines. Even though our current set (350 target proteins) is smaller than the existing sets, our set contains 239 highly expressed RBPs, of which 93 were uniquely present in our set (Figure 5F).

In addition, our database allows users to browse the essential information to start tagging experiments including pre-designed gRNA sequences, pLDDT scores of regions surrounding a target intron, target expression levels, and Venus-positivity rates after transfection displayed on the genome browser, sortable/searchable tables or individual web pages. Moreover, the microscopy and Western blotting results are continually being updated. The microscopy and western blotting results will be updated once the data are obtained. The availability of individually isolated plasmids that could be used for intronic NHEJ knock-in experiments in different human cell lines using different epitope tags makes our gRNA set a unique resource.

## Data availability

The database is available at <https://rnabio.naist.jp/atlas/>.

## Funding

Y.H. was supported by the research fellowship from the NAIST for the Creation of Innovation in Science and Technology, and the JSPS Research Fellowship for Young Scientists (Grant-in-Aid for JSPS Fellows, 24KJ1692). N.Y. and K.M were supported by the NAIST Granite Program. Works at K.O.’s lab was supported by the JSPS Found for the Promotion of Joint International Research (Home-Returning Research Development Research, 17K20145), Academic Assistant Grant by Office for Gender Equality at NAIST and the Takeda Science Foundation. T.K.’s lab was supported by JSPS KAKENHI (Grant-in-Aid for Scientific Research (B) 20H03468). N.K. was supported by JSPS KAKENHI (Grant-in-Aid for Early-Career Scientists 23K14546).

## Supplementary data

Supplementary data is available at NAR online.

## Conflict of interest

The authors declare that they have no conflicts of interest with the contents of this article.

## Acknowledgements

The authors thank Hiroshi Ito for the Venus plasmid and the Life Science Collaboration Center (LiSCo) facilities for microscopy. We also thank Masami Shiimori, Ayako Isotani, Toshiaki Shigeoka, Yasumasa Ishida, Minato Hirano and Kentaro Yoshii for constructive feedback on the manuscript and the website. We also thank Yae Toda, Junko Iida, Sojiro Ukeda, Masayoshi Shiina, Riku Nakao and Yuto Shimoda for technical assistance.

## Author contributions

Y.H., P.L.L.H., Y.S., and K.O. writing-review and editing; Y.H. and K.O. writing-original draft; Y.H. and K.O. visualization; Y.H. and K.O. software; Y.H., P.L.L.H., Y.S., N.Y., S.M., Y.S., N.K., M.K. and K.M. resources; Y.H., P.L.L.H., Y.S., N.Y., S.M., Y.S., N.K., T.K., and K.O. methodology; Y.H. and K.O. investigation; N.K., T.K., Y.H. and K.O. formal analysis; N.K., T.K., Y.H. and K.O. funding acquisition; T. K. and K.O. supervision; T.K. and K.O. project administration; K.O. conceptualization.

---

**Figure S1.**
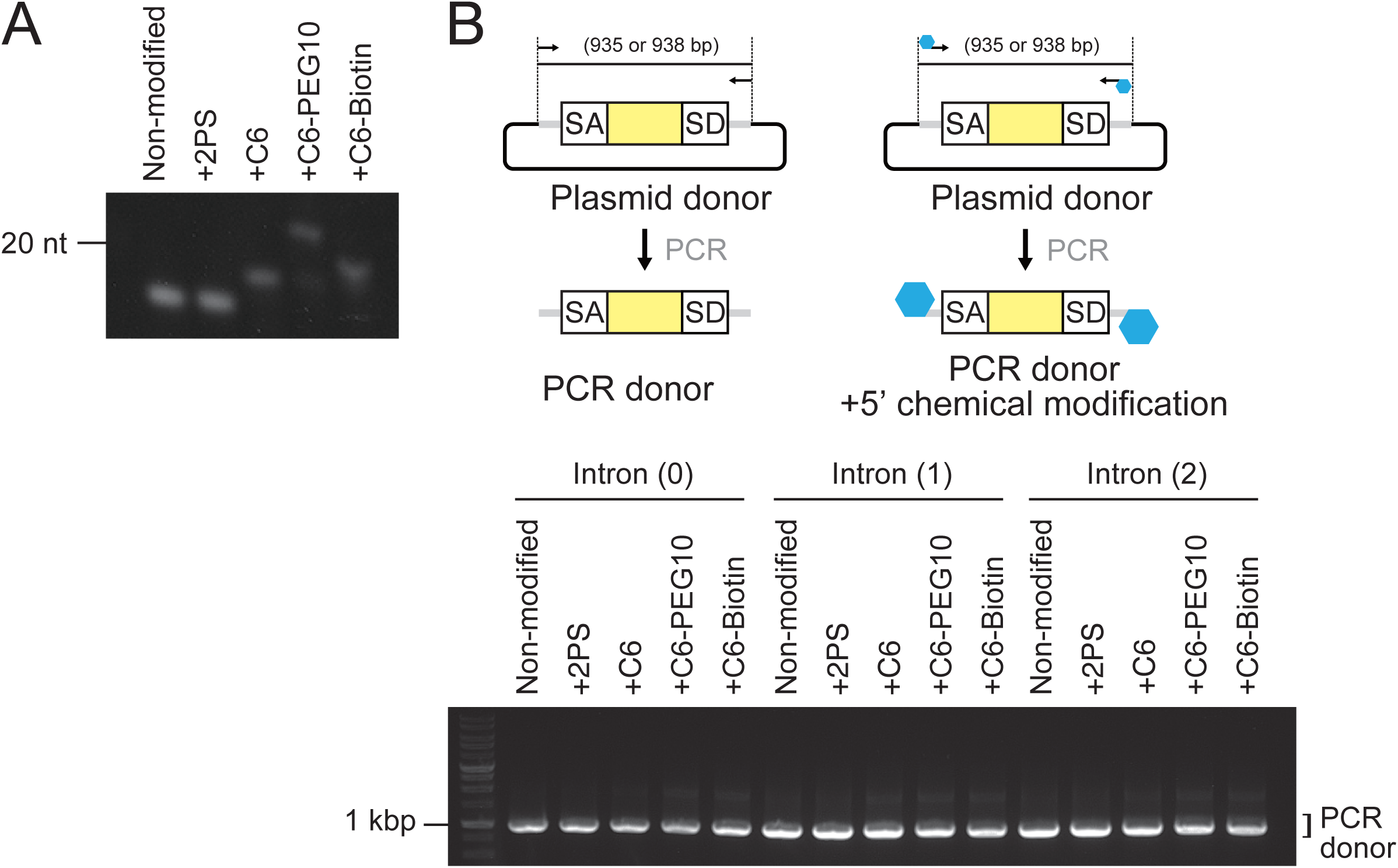
Verification of chemical modified PCR products. (A) Representative urea-gel picture of modified primers. 10 pmol primers were resolved using 15% urea-gel and stained with SYBR Gold. (B) Representative agarose gel picture of the PCR knock-in donors with 3 different chemical modifications.

**Figure S2.**
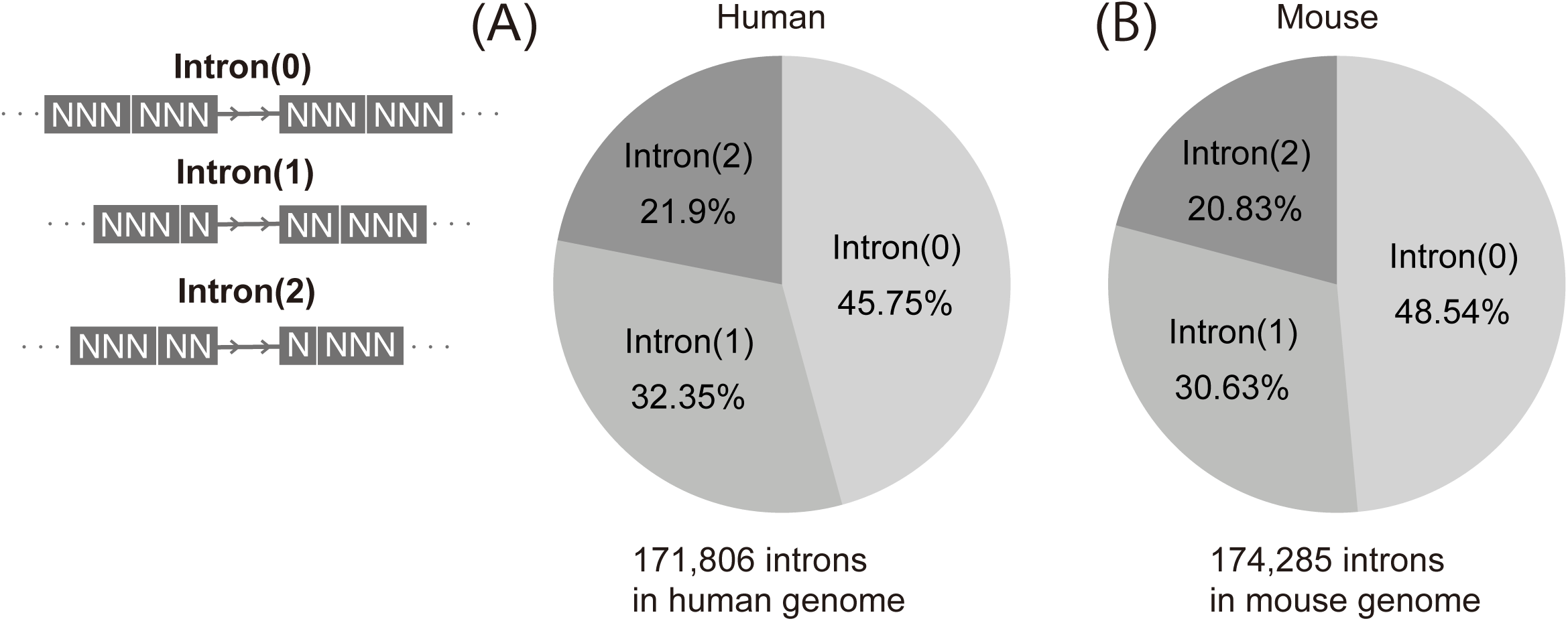
Percentages of intron phases in CDSs in the human and mouse genomes. The pie chart shows the fraction of Intron (0), (1) and (2) among the 171, 806 introns in the human 16, 993 genes (A) and the 174, 285 introns in the mouse 17, 715 genes (B).

**Figure S3.**
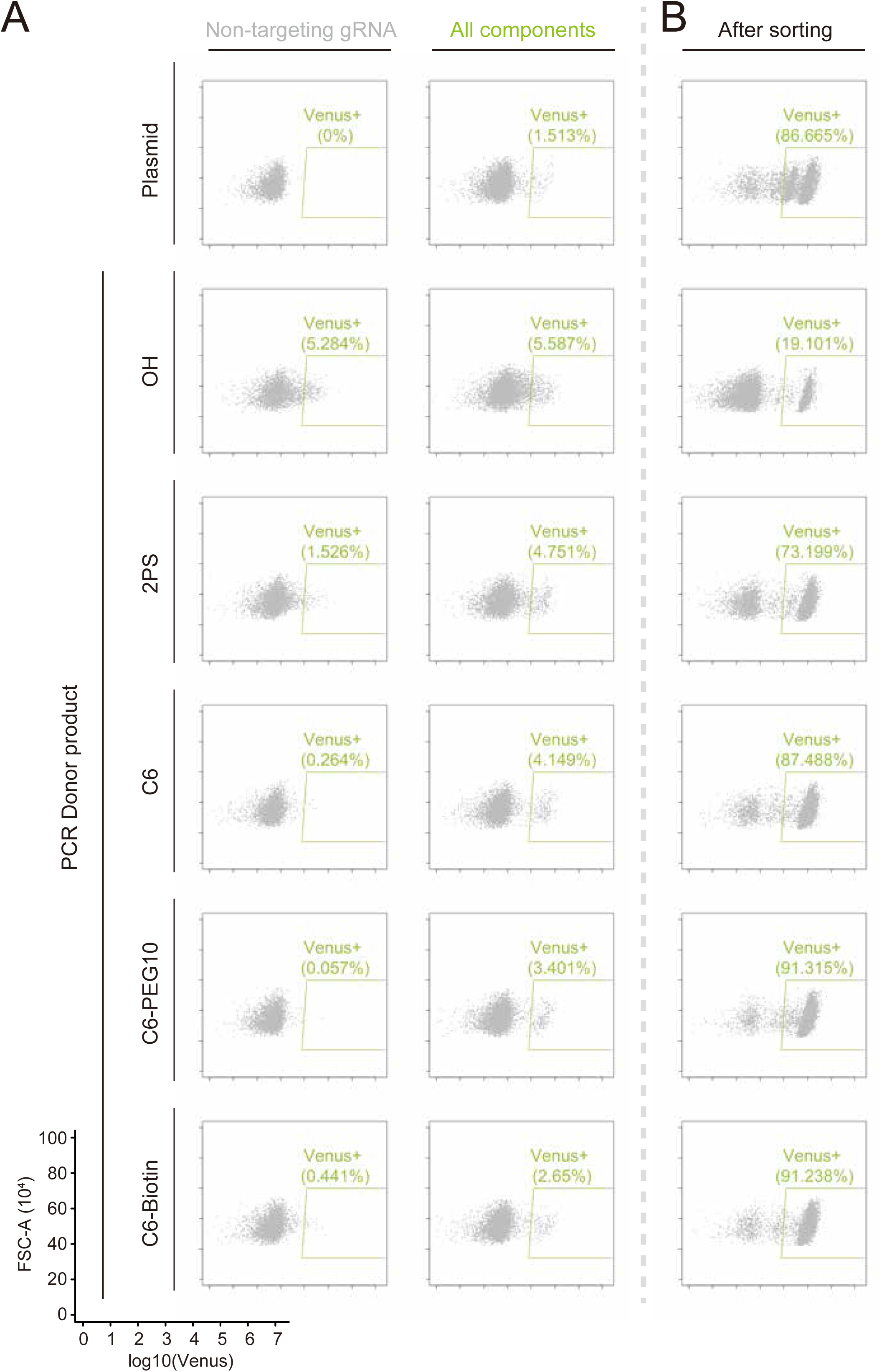
Fluorescence intensity of cells after Venus knock-in at HNRNPA1-Intron 1. Scatter plots of log10-transformed Venus intensity (x-axis) and FSC-A (y-axis) are shown. Flow cytometry analysis was done 2 days after transfection (A) or 1 week after sorting (B).

**Figure S4.**
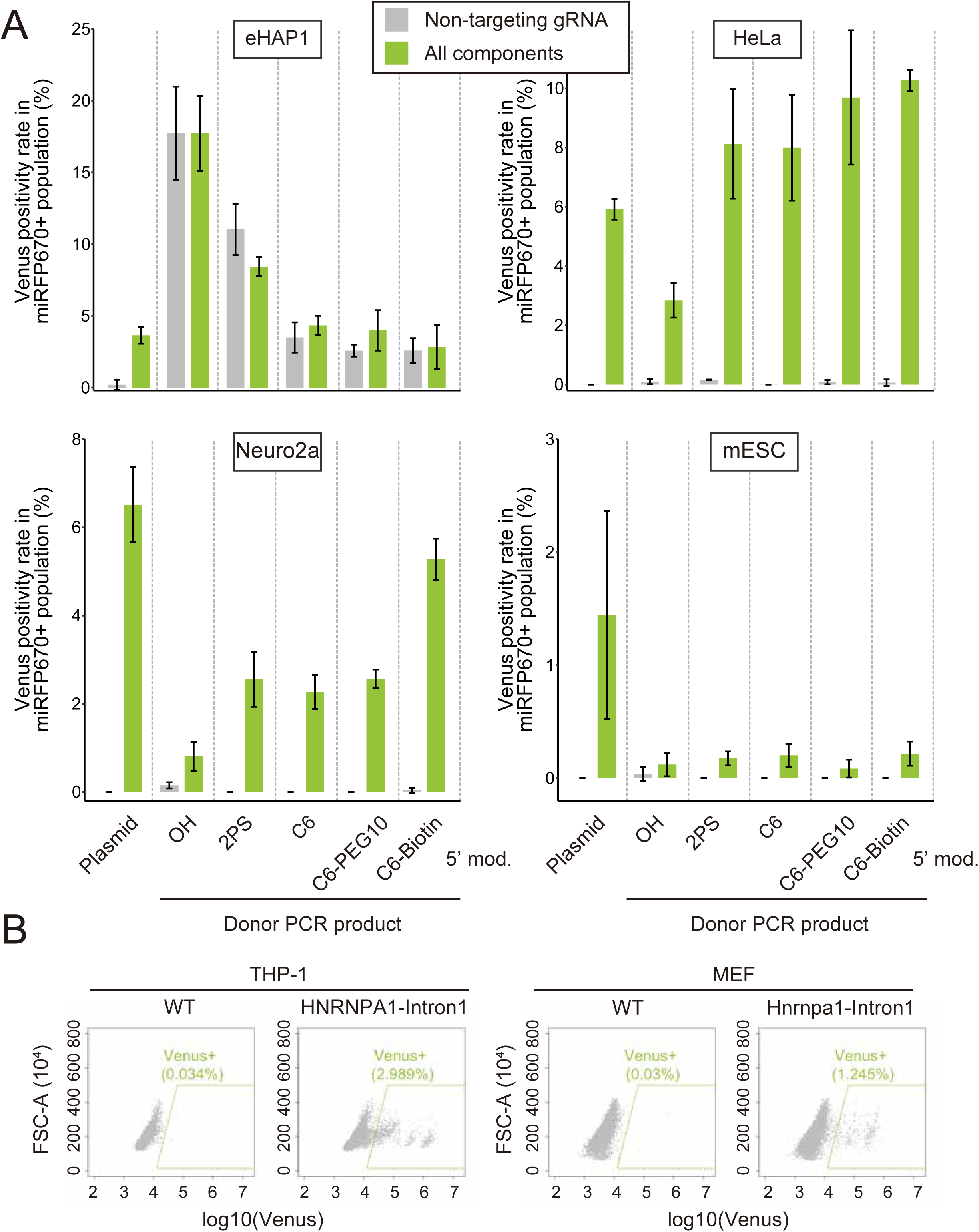
Venus positivity rates by various donors. (A) Flow cytometry data of knock-in cells 2 days after transfection. Grey bars show the results with the non-targeting gRNA expression plasmid, gRNA expression plasmid against the donor sequence and the Venus donor (in the case of plasmids) or the PCR product. Green bars show the results with the complete set gRNA expression plasmid and donor DNA. (B) Flow cytometry data of knock-in THP-1 and MEF cells 1-2 weeks after transfection. Scatter plots of log10-transformed Venus intensity (x-axis) and FCS-A (y-axis) are shown.

**Figure S5.**
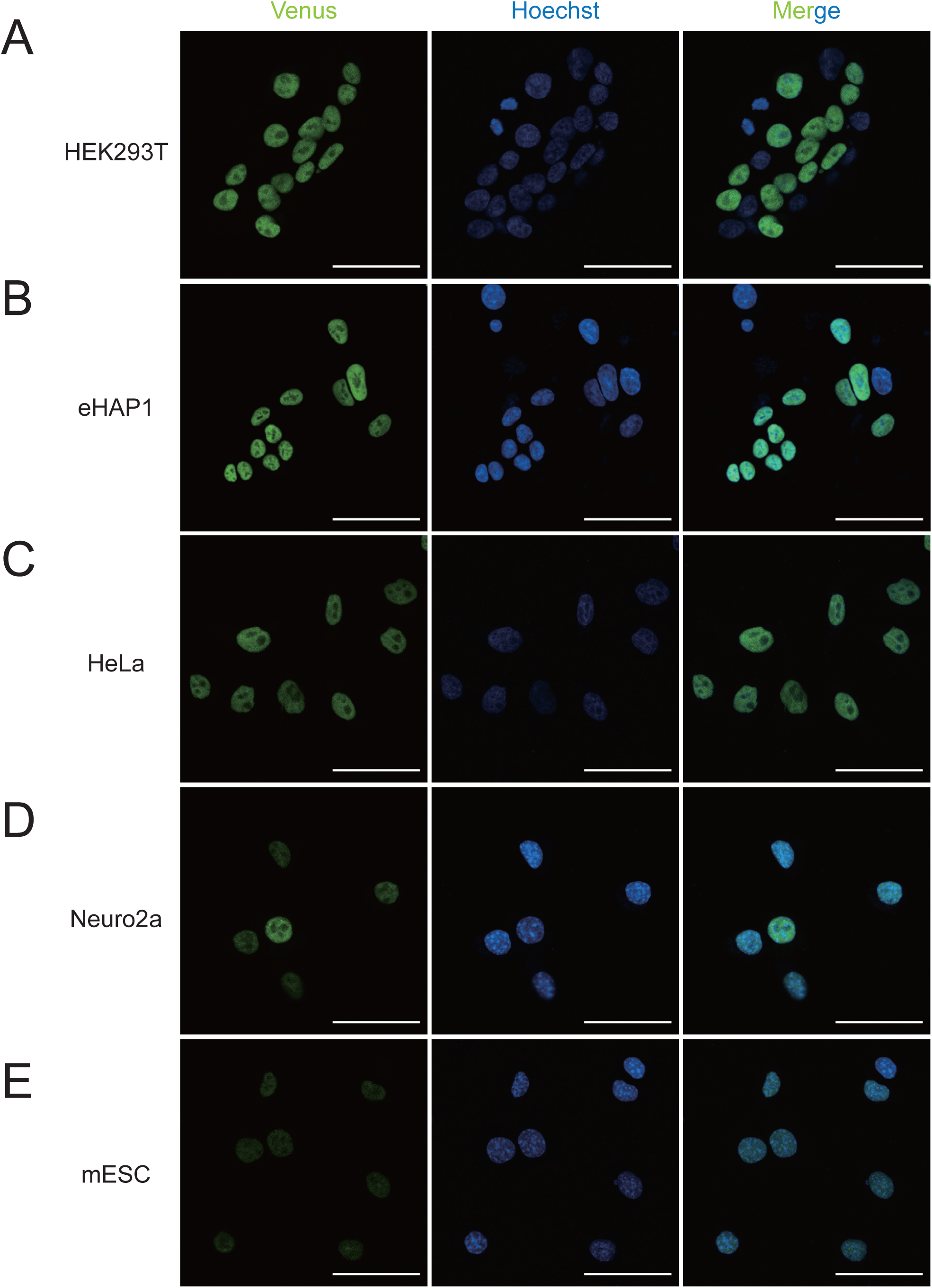
Subcellular localization of HNRNPA1-Venus. The fusion protein localization of HNRNPA1-Venus was observed in the bulk knock-in HEK293T (A), eHAP1 (B), HeLa (C), Neuro2a (D) and mESC (E) with a 20× objective lens, and scale bars correspond to 50 μm. PCR product with 5’-C6 modification were used except for mESC (plasmid donor was used in the case of mESC).

**Figure S6.**
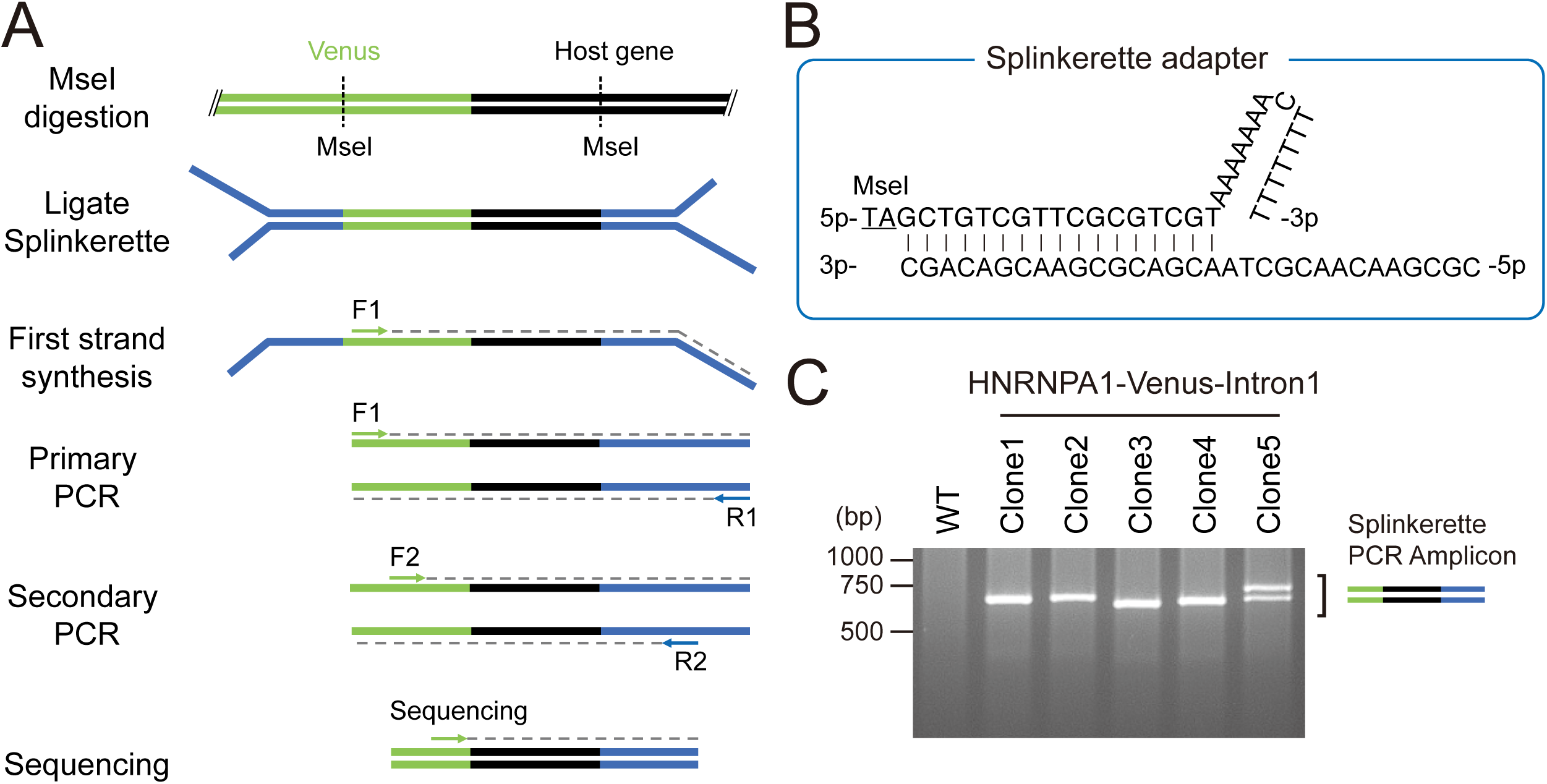
Determination of insertion sites using splinkerette PCR. (A) Schematic of the splinkerette PCR method. Genomic DNA digested with Mse I is ligated with splinkerette adapter and DNA fragments were amplified using Venus-specific primer and splinkerette adapter-specific primer. (B) The sequences and the predicted structure of the Splinkerette adapter. The upper strand has the TA sticky 5’ overhang (underline). (C) Representative agarose gel picture of splinkerette PCR using the HNRNPA1 knock-in eHAP1 clones derived from single cells. Most clones produced single bands at the expected around 700 bp size, except clone 5 that gave rise to two bands.

**Figure S7.**
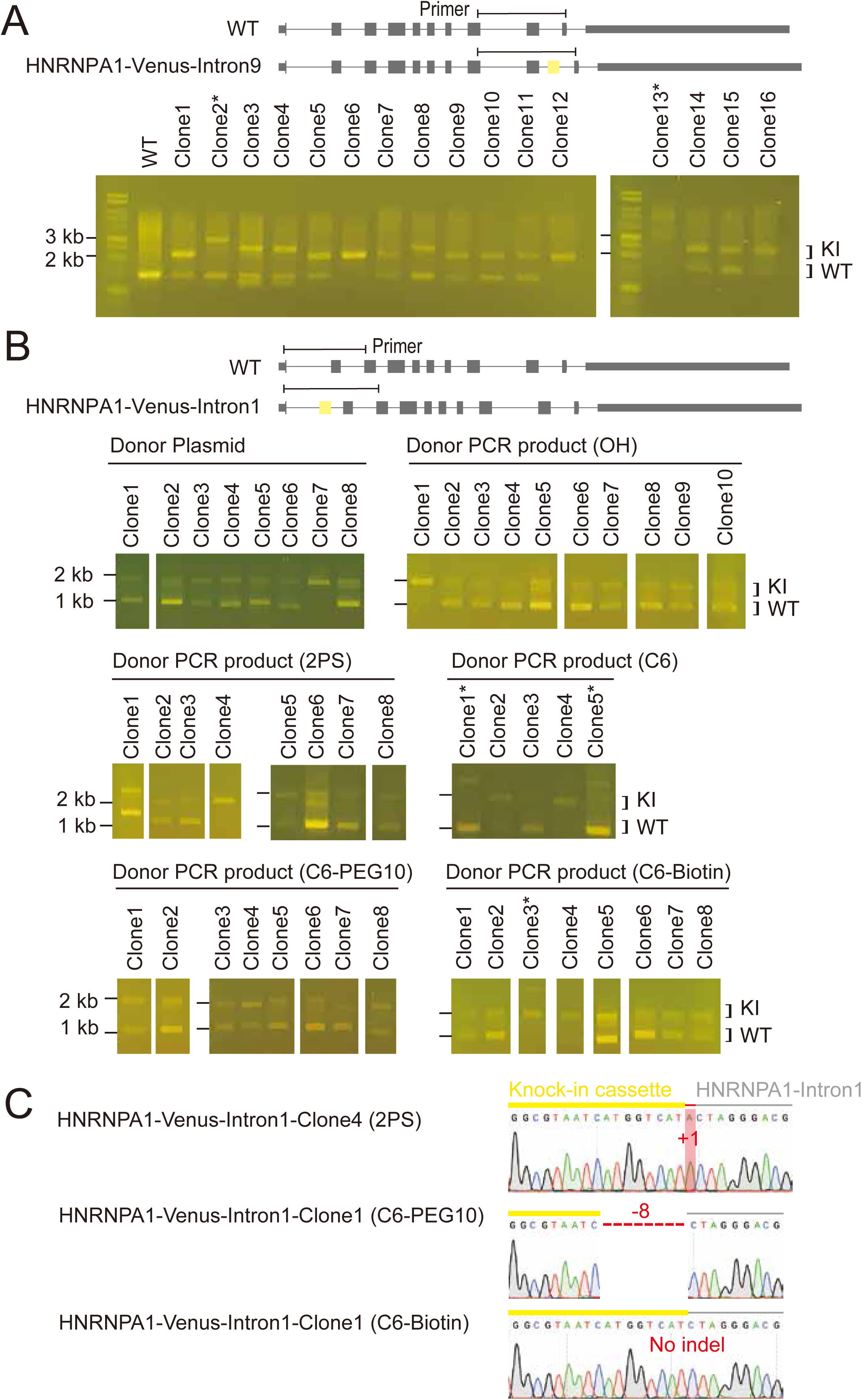

**Figure S8.**
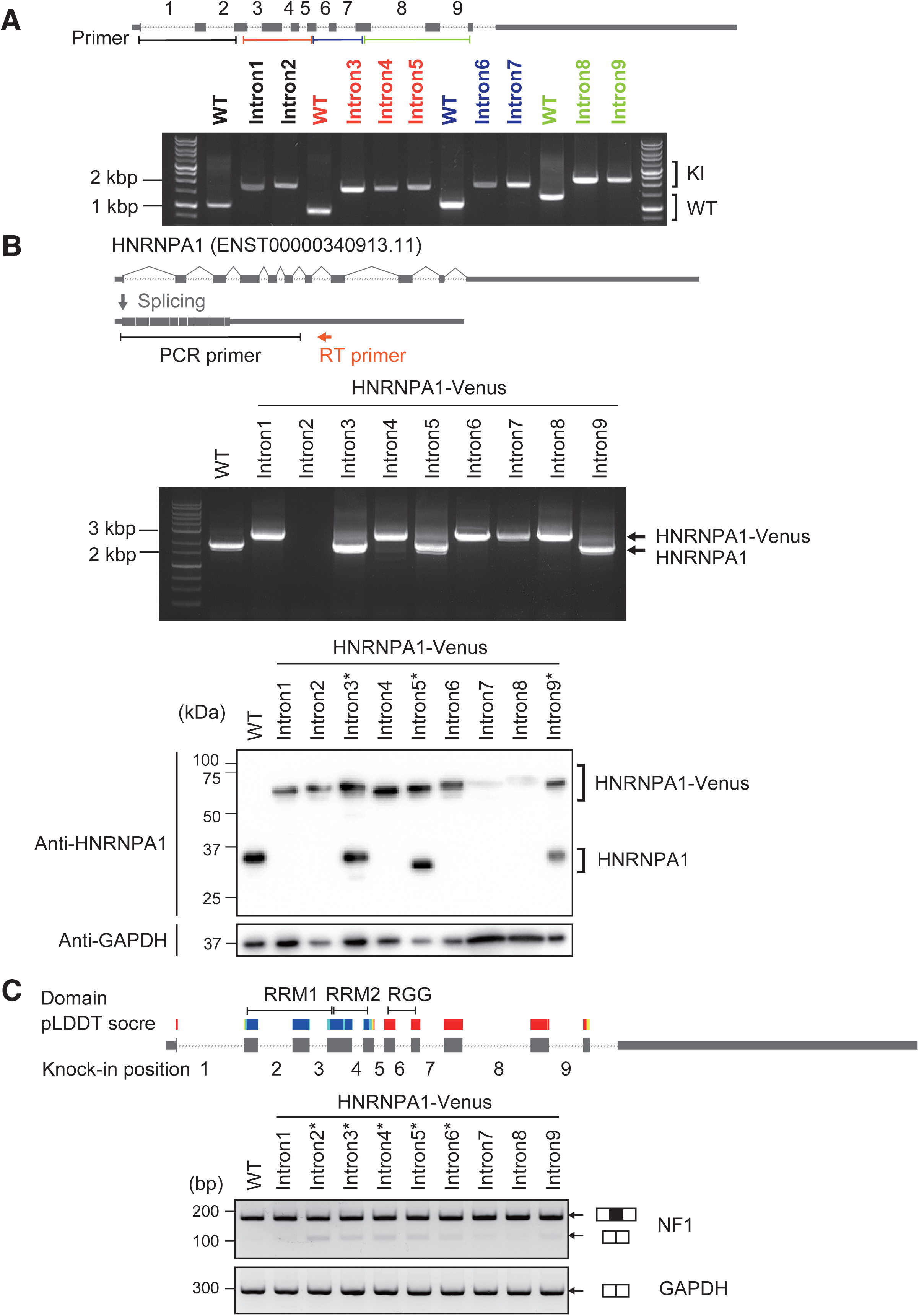

**Figure S9.**
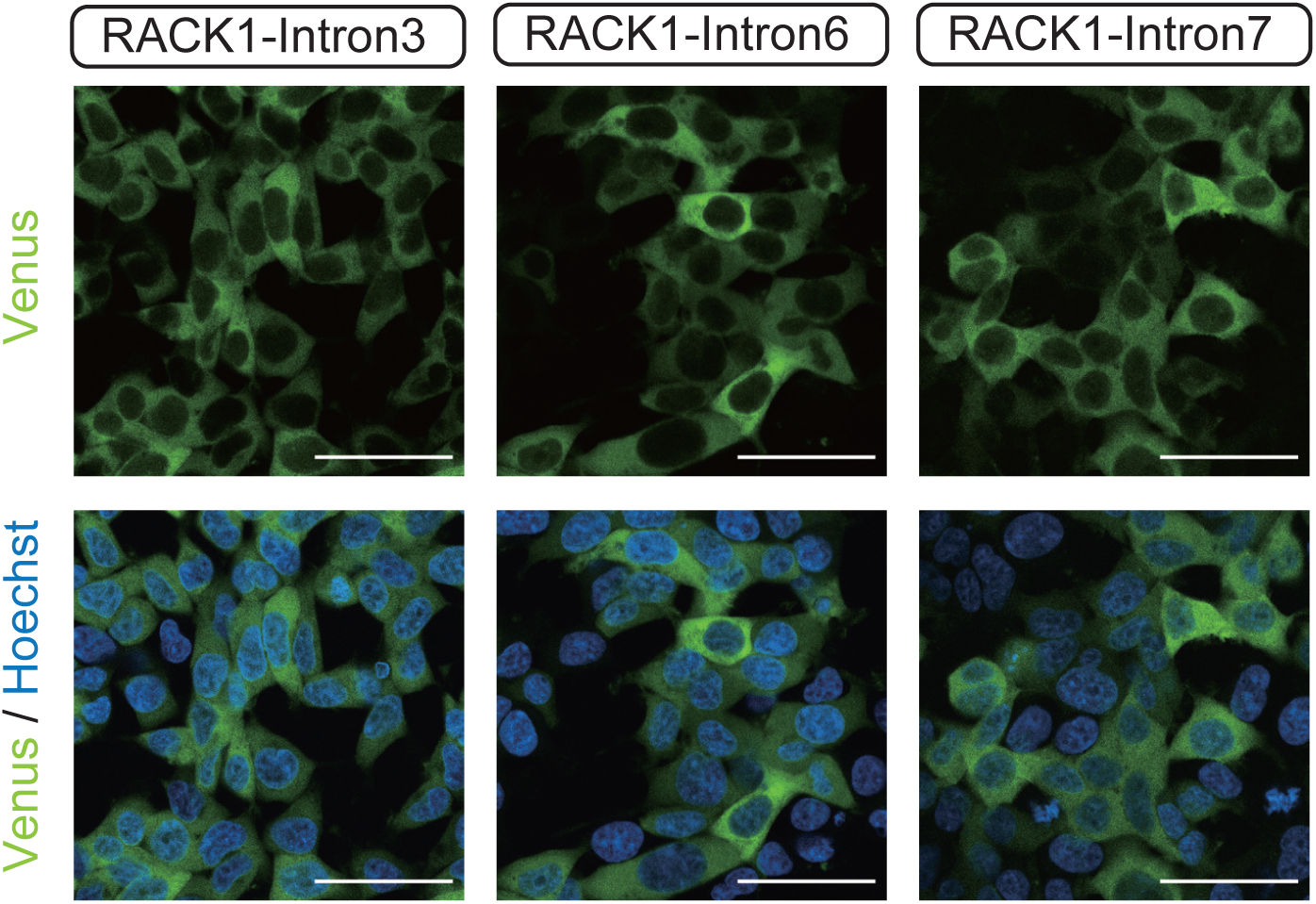
Subcellular localization of RACK1 fusion proteins. The fusion protein localization was analyzed in RACK1-mVenus-Intron 3, 6 and 7 bulk cell population with a 20× objective lens. Scale bars correspond to 50 μm.

**Figure S10.**
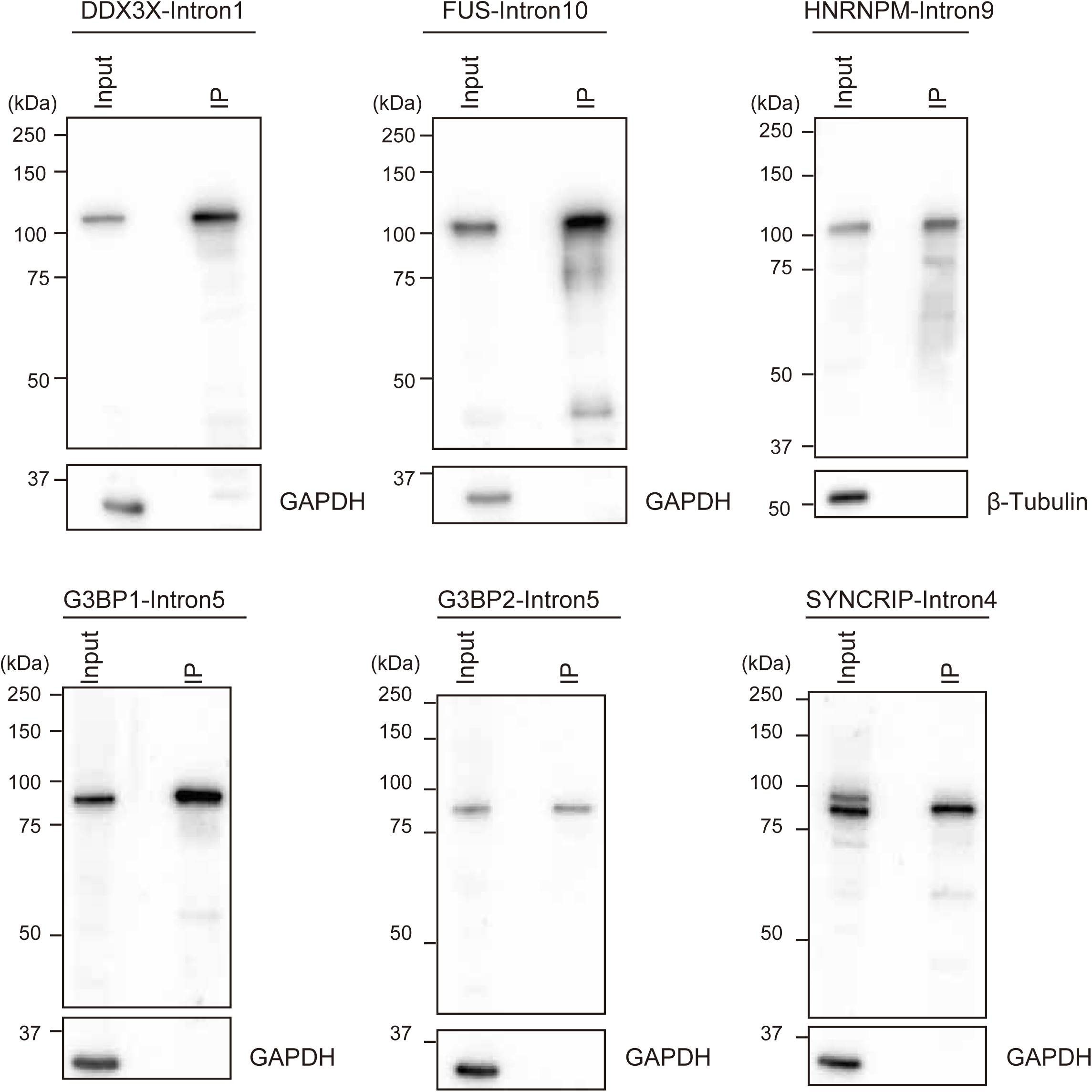
Summary of immunoprecipitations. Western blot of mVenus-tagged protein after immunoprecipitation using anti-GFP VHH antibody. The FUS-Intron10, DDX3X-Intron1, G3BP1-Intron5, G3BP2-Intron5, HNRNPM-Intron9 and SYNCRIP-Intron4 lines derived from single cells were immunoprecipitated with the anti-GFP VHH antibody. Samples were resolved 8% SDS-PAGE and bands were detected by an anti-GFP and anti-β-Tubulin or anti-GAPDH antibodies sequentially.

## Notes

### Competing Interest Statement

The authors have declared no competing interest.

### Summary of Updates

Total number of genes increased to 350; Figure S7 and Figure S9 added; authors added

